# Identifying sensor species to predict critical transitions in complex ecosystems

**DOI:** 10.1101/472878

**Authors:** Andrea Aparicio, Jorge X. Velasco, Claude H. Moog, Yang-Yu Liu, Marco Tulio Angulo

## Abstract

Ecosystems provide key services needed for human well-being, such as purifying air and regulating our health, many of which are disrupted when these ecosystems undergo a critical transition to undesired states. Detecting early-warning signals of these critical transitions remains challenging for complex ecosystems with a large number of species. Here we built a mathematical formalism to identify minimal sets of “sensor species” from which we can determine the state of a whole ecosystem, allowing us to predict a critical transition in an ecosystem by monitoring a minimal subset of its species. We rigorously prove that minimal sets of sensor species can be generically identified knowing only the structure of the ecological network underlying the ecosystem, regardless of its population dynamics. We numerically validated our formalism to predict critical transitions in large complex ecosystems, and then we applied it to experimental data of a critical transition in a lake food-web. Our results contribute to better monitoring complex ecosystems, especially those with poorly known population dynamics such as host-associated microbial communities.

## INTRODUCTION

Ecosystems provide many services that we humans need for our well-being [1], from supplying food and purifying air and water [2], to regulating our health via the microbial communities in and on our bodies [3]. An ecosystem can undergo “<u>critical transitions</u>” (i.e., abrupt changes) to undesirable states where the services it provides decrease, some of its species become extinct, or the complete ecosystem collapses [4]. For example, fisheries can transition to states of low productivity [5], healthy coral-dominated reefs can transitions to unhealthy algal-dominated states [6], and rangelands can transition to low productivity states dominated by shrubs with little grass [7]. Detecting early-warning signals of such critical transitions is a necessary step to prevent them [8, 9]. Several methods have been developed for detecting early-warning signals of critical transitions using time-series of the ecosystem [10], including detecting a so-called “critical slowing down” of the system’ response reflected by an increase in its variance [11] or autocorrelation [12]. Detecting these early-warning signals for large complex ecosystems remains a major challenge due to the complex population dynamics they may exhibit, together with the large number of involved species [4, 13]. Actually, because in some cases it is simply unfeasible to measure the activity of all the species involved in the ecosystem, critical transitions may remain totally undetected since their early-warning signals can be weakly present [14] or totally absent [15] in the species that are measured.

To overcome the above challenge, here we identify minimal sets of “<u>sensor species</u>” for an ecosystem, defined as a subset of species that determines the whole state of the ecosystem. We show that the sensor speciescanbeusedto detect early-warning signals of critical transitions happening in any part of the ecosystem. To identify the sensor species of an ecosystem, we developed a mathematical formalism based on the new notion of <u>structural observability</u> for nonlinear systems, which generalizes to uncertain nonlinear systems the notion of local observability introduced in nonlinear control theory [16]. The notion of structural observability we propose is also a nonlinear generalization of the notion of “linear structural observability” developed for linear systems, which as been extensively studied in recent years from the perspective of Network Science (see, e.g., [17]). A system is called structurally observable from a set of measured variables if, for almost all dynamics that the system can have (linear or nonlinear), these measured variables locally determine the complete state of the system. With this notion, a set of sensor species can be characterized as a subset of species that render the ecosystem structurally observable. Our main theoretical result shows that the structural observability of an ecosystem depends only on the topology of its ecological network, regardless of its particular population dynamics. This result enabled us to construct an efficient algorithm to find a minimal set of sensor species in large ecosystems knowing only the structure of their underlying ecological network. We then constructed early-warning signals based on sensor species, numerically validating their performance on large ecosystems. In particular, our validation shows that sensor species provide early-warning signals that are robust to errors in the ecological network used to identify them. Finally, we identified the sensor species from an experimental lake food-web, and then used experimental data to verify that this sensor species predicts a critical transition. Overall, our results illustrate how embracing the underlying complexity of ecological networks can improve our ability to monitor large ecosystems and detect critical transitions before they occur.

## IDENTIFYING SENSOR SPECIES

We consider an ecosystem of *N* species labeled as X = {1, …, *N*} whose <u>state</u> *x*(*t*) = (*x*_1_(*t*), …, *x_N_*(*t*)) ∈ ℝ^*N*^ at time *t* is given by the abundance of all its species. Here *x_i_*(*t*) denotes the abundance of the *i*-th species at time *t*. Our goal is to identify a minimal subset Y of species to measure Y ⊆ X, which we call <u>sensor species</u>, such that their abundance determines the abundance of the rest of the species X\Y of the ecosystem.

### What makes a sensor species?

Consider the ecological network 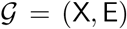 underlying the ecosystem, where the set of nodes X corresponds to the species, and the directed edge (*j → i*) ∈ E represents a direct ecological interaction from the *j*-th species to the *i*-th one (i.e., a direct promotion or inhibition of growth). This ecological network serves as the conduit through which changes in the abundance of some species are propagated to otherspecies inthe ecosystem. Therefore, to have a valid set Y of sensor species, a necessary condition is that G contains a path that propagates a change in any unmeasured species to some measured species (i.e., a path from any species in X \ Y to some species in Y). To understand if such condition is not only necessary but also sufficient to obtain a valid set of sensor species, consider the toy three-species ecosystem shown in Fig. 1A. According to the above argument, any of its three species should be a valid sensor species because there is a path between any of them. We tested this hypothesis choosing Y = {1} as a solo sensor species and simulating three population dynamics models for this ecosystem having increasing complexity (Figs. 1B to D). The simplest (and less realistic) population dynamics that we consider is the linear dynamics

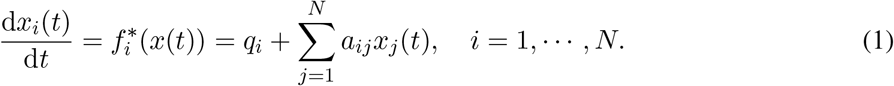

**FIG. 1:**
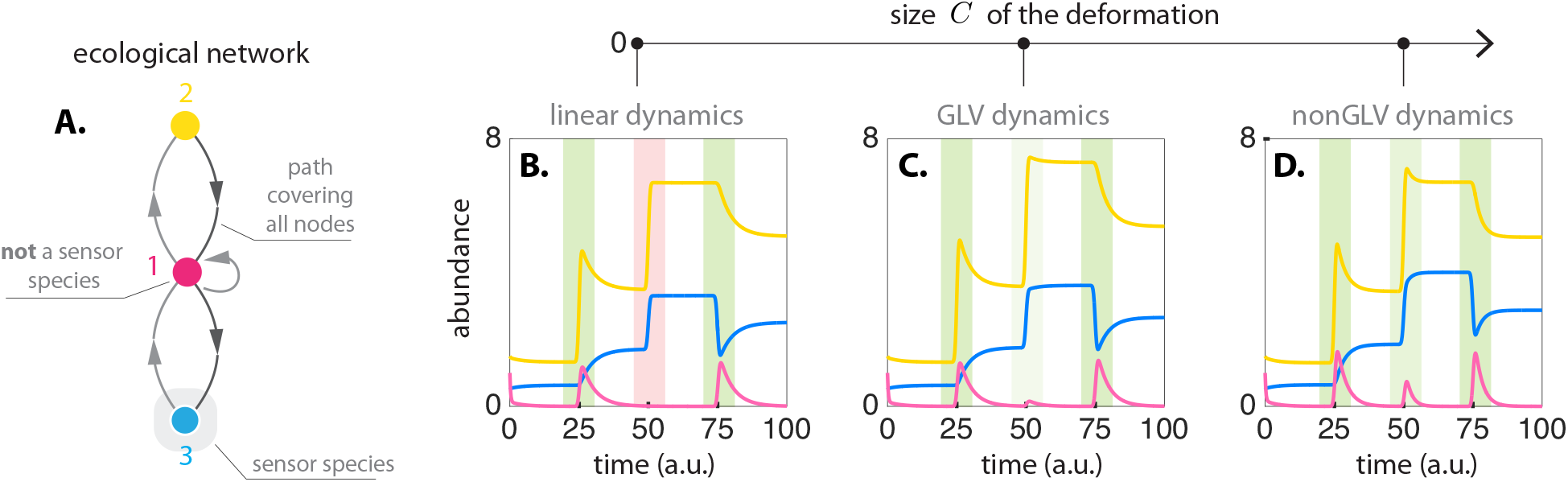
Sensor species depend on the ecosystem’s ecological network and its population dynamics. **A.** Toy ecological network with *N* = 3 species X = {1,2,3}. **B.** Abundance of species using the linear population dynamics *ẋ*_1_ = −4*x*_i_ + 1.6(*x*_2_ − 2*x*_3_) − 0.1, *ẋ*_2_ = −0.2*x*_1_, *ẋ*_3_ = 0.15*x*_1_. Perturbations are applied to change the abundance of species 2 at *t* = {25, 50}, and to change the abundance of species 3 at *t* = {50, 75}, see color stripes. The changes in species 2 or species 3 are detected by a change in species 1 (green stripes). However, the simultaneous change in species 2 and 3 at t = 50 remains undetected in species 1 (red stripe), implying that with linear dynamics, species 1 cannot be a sensor species. This is mathematically characterized by the lack of observability of the linear population dynamics with respect to species 1. **C.** Abundance of species using the GLV dynamics *ẋ*_1_ = (1 − *θ*_1_ + *θ*_1_*x*1)(−4*x*1 + 1.6(*x*_2_ − 2*x*3) − 0.1), *ẋ*_2_ = (1 − *θ*_1_ + *θ*_1_*x*_2_)(−0.2*x*_1_), *ẋ*_3_ = (1 − *θ*_1_ + *θ*_1_*x*_3_)(0.15*x*_1_), and *θ*_1_ = 0.05. Note this corresponds to a deformation with size *C* = 1. The simultaneous change at t = 50 is now detected by a change in species 1, making species 1 a sensor species (light green stripe). Mathematically, this is characterized by the local observability ofthe GLV dynamics with respect to species 1 (Example 2 in Supplementary Note 1.2). Note, however, that the response of species 1 to the simultaneous change is very small compared to its response to individual changes. **D.** Abundance of species using a deformation of size *C* = 2 as population dynamics. Its equations are *ẋ*_1_ = (1 − *θ*_1_ + *θ*_1_*x*_1_)(−4*x*_1_/(1 + *θ*_2_*x*_1_) + 1.6(*x*_2_/(1 + *θ*_2_*x*_2_) − 2*x*_3_/(1 + *θ*_2_*x*_3_)) − 0.1), *ẋ*_2_ = (1 −*θ*_1_ + *θ*_1_*x*_2_)(−0.2*x*_1_/(1 + *θ*_2_*x*_1_)), *ẋ*_3_ = (1 −*θ*_1_ + *θ*_1_*x*_3_)(0.15*x*_1_/(1 + *θ*_2_*x*_1_)) with *θ*_1_ = 0.05 and *θ*_2_ = −0.05. The simultaneous change at *t* = 50 is also detected by a change in species 1, making species 1 a sensor species (green stripe). The response of species 1 is largerthan in Panel C.

Here, *q* = (*q_i_*) ∈ ℝ^*N*^ are migration rates from/to neighboring ecosystems, and *A* = (*a_ij_*) ∈ ℝ^N×N^ is the interaction matrix of the ecosystem. Namely, the parameter aij quantifies the direct interaction from the *j*-th species to the *i*-th one. A linear model is useful if the ecosystem remain close to an equilibrium point, and it is also the simplest dynamics that could approximate an ecosystem. Under the linear dynamics of Eq. (1), we find that a change in the abundance of either species 2 or 3 is always detected by a change in the abundance of the species 1, as desired in a sensor species (green stripes in Fig. 1B). However, a simultaneous change in the abundance of species 2 and 3 can remain completely undetected by species 1 (red stripe in Fig. 1B). This means that Y = {1} cannot be a sensor species for this ecosystem under linear dynamics. As the dynamics of the ecosystem become more complex, a simultaneous change in specie 2 and 3 can be detected in species 1 (Figs. 1C and D). This implies that more complex population dynamics can render Y = {1} a valid sensor species.

The reason why species 1 alone cannot be a sensor species for the linear population dynamics of Eq. (1) is because there are two temporal responses of the ecosystem —without perturbations, and with simultaneous perturbations to species 2 and 3— that give the exact same response in species 1. This makes it impossible to decide which of the two responses is actually occurring in the ecosystem from measuring species 1 alone. Therefore, if we denote by *y*(*t*) ∈ ℝ^|Y|^ the abundance of the species in a valid set Y of sensor species, for each response of the sensor species {*y*(*t*), *t* ≥ 0} there should be one and only one response of the whole ecosystem {*x*(*t*),*t* ≥ 0}. This condition defines the notion of <u>observability</u> of a dynamic system with respect to the measured variable *y*, a cornerstone concept in modern control theory [16]. This observation motivated us to <u>define a set of sensor species</u> as a set of species that render the ecosystem (locally) observable from them. Species 1 in Fig. 1A fails to be a solo sensor species because it does not render this ecosystem with linear dynamics observable (Example 1 in Supplementary Note 1.2). By contrast, species 2 or 3 in Fig. 1A render this ecosystem observable (Examples 2 and 3 in Supplementary Note 1.2), making them valid sensor species.

Mathematically, the local observability of a system is defined as the existence of a locally invertible map (called the <u>observability map</u>) that allows writing locally the full state of the ecosystem *x*(*t*) as a function of the sensor species and its time derivatives {*y*(*t*), *ẏ*(*t*), … *y*^(*N_−p_*)^(*t*)}. Here *p* = |Y| is the number of sensor species (see Definition 2 in Supplementary Note 1.2 for details). However, notice that the observability map not only depends on the ecological network of the ecosystem, but it also depends on its population dynamics (including its particular parameters). This means that the same ecological network can be observable for some population dynamics, and unobservable for other population dynamics. For example, in the ecological network of Fig. 1A, using the Generalized Lotka-Volterra (GLV) as model for its population dynamics makes the ecosystem observable with respect to species 1, because this species can detect a simultaneous change in species 2 and 3 (Fig. 1C). That is, for this ecological network Y = {1} is a solo sensor species for GLV dynamics, but not for linear dynamics.

In summary, the above toy example illustrates two important aspects that must be considered when characterizing the sensor species of an ecosystem. First, it might not be sufficient to choose as sensor species a set of species that “receives” information from all other species (e.g., all species in the lowest trophic level). Second, in general, the set of sensor species of an ecosystem does not only depend on its ecological network but also on its population dynamics. This second point may suggest that it is impossible to identify the sensor species of most ecosystems because of the very limited knowledge we have of their population dynamics. Indeed, while it is feasible to map the ecological networks underlying a wide class of ecosystems, from food-webs [19] to microbial communities [20], it remains much more challenging to infer their population dynamics [21, 22]. In what follows we build a mathematical formalism to circumvent this seemingly unavoidable challenge, letting us identify the sensor species of an ecosystem only from the topology of its underlying ecological network.

### Characterizing minimal sets of sensor species from ecological networks

Knowing only the ecological network 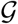 ofanecosystem, weareledto analyze the observability of the class 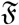 of all population dynamics that such ecosystem may have. In general, the population dynamics of an ecosystem takes the form of a set of *N* coupled nonlinear differential equations

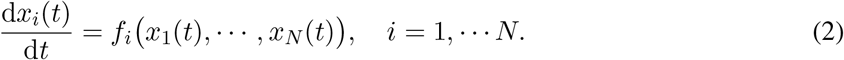

Here, the function *f* = (*f*_1_, …, *f_N_*)^*T*^: ℝ^*N*^ → ℝ^*N*^ models the intra- and inter-species interactions in the ecosystem. Different population dynamics in the ecosystem correspond to different *f*’s, meaning that its functional form can significantly change between ecosystems. To acknowledge this situation, we consider that *f* is some unknown <u>meromorphic function</u> —each of its entries is the quotient of analytical functions of *x*. This assumption is very mild in the sense that it is satisfied for most population dynamics models [21].

To construct the class 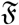, we first define the network or <u>graph</u> 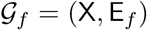 associated with a meromorphic function *f* as follows: the edge (*j → i*) ∈ E_*f*_ exists if *∂f_i_/∂x_j_* ≢ 0 (see details in Definition 3 in Supplementary Note 1.3). Note that (*j → i*) ∈ E_*f*_ simply means that *ẋ_i_* directly depends on *x_j_*. Next, define the set of <u>base models</u> of the ecosystem, representing the simplest population dynamics that the ecosystem could take. We thus chose the base models as all linear models *f*\* of Eq. (1) such that 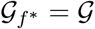. Finally, we construct the class 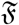 as containing all base models, together with all <u>deformations</u> of each base model.

A function *f*(*x*) is a deformation of *f*^*^(*x*) if: (i) both have the same graph (i.e., 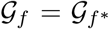); (ii) there exists a meromorphic function *f̃*(*x*; *θ*): ℝ^*N*^ × ℝ^*C*^ → ℝ^*N*^ such that *f̃*(*x; θ*) = *f*(*x*); and (iii) the identity *f*(*x; θ*) = *f*\*(*x*) holds. We call the minimal integer *C* ≥ 0 that satisfies these conditions the <u>size</u> of the deformation. A rather general class of population dynamics can be described by deformations of Eq. (1), including for example

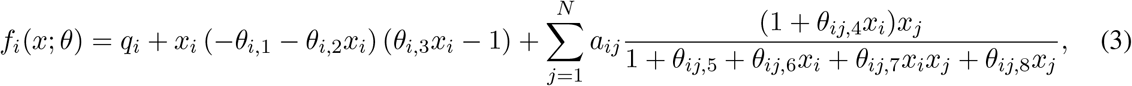

for *i* = 1, …, *N*. Here, *θ*_*i*,1_ are intrinsic growth rates, 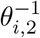 are the carrying capacities of the environment, 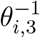 are the Allee constants, and the rest of the parameters 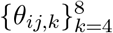 characterize the saturation of the functional responses [21]. Other population dynamics having so-called “higher-order” interactions 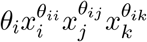 can also be described as deformations. The notions of base models and their deformation was recently used to identify the “driver species” that allows controlling a microbial community [23]. Here, we are adapting these notions for characterizing the “dual” concept of sensor species.

We call 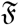 <u>structurally observable</u> with respect to a subset of species Y ⊆ X if almost all of its base models and almost all of their deformations are locally observable with respect to *y* ∈ Y. In other words, except for a zero measure set of “singularities” in the population dynamics that the ecosystem may have, the species in Y is a valid set of sensor species. This formulation allowed us to disentangle the relation between the population dynamics of an ecosystem, its ecological network, and its sensor species. First, we prove that increasing the size of a deformation cannot generically invalidate a set of sensor species (Proposition 1 in Supplementary Note 1.3). This result reduces the analysis to the subset of deformations in 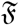 with size *C* = 0. Our second result proves that 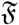 is structurally observable if and only if the network 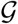 satisfies the following two conditions: (i) there is a path from each node in X to a node in Y; and (ii) there is a disjoint union of cycles and paths that covers X (Theorem 3 in Supplementary Note 1.3). Recall that a path in 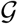 is a sequence of consecutive edges (e.g., {1 → 2 → 3}), and a cycle is a path that starts and end in the same node (e.g., {1 → 2 → 1}). A set of edges covers X if each of its nodes is part of some edge. The conditions (i) and (ii) above allowed us to map the problem of finding minimal sets of sensor species to that of solving a maximum matching problem in 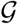 (Supplementary Note 1.4). Such a problem can be solved in polynomial time, letting us efficiently find minimal sets of sensor species for ecosystems with a large number of species.

## CONSTRUCTING EARLY-WARNING SIGNALS FROM SENSOR SPECIES

Let Y ⊆ X be a set of p sensor species of the ecosystem, and denote by *y*(*t*) ∈ ℝ^*p*^ the abundance of those sensor species at time *t*. For a <u>lag size</u> *L* > 0, define *Y_L_*(*t*) = [*y*(*t*),*y*(*t − L*), *y*(*t* − 2*L*), …, *y*(*t* − (*N* − *p*)*L*)]^*T*^ as the <u>sensor space</u> vector, consisting of the abundance of the sensor species and (*N − p*) of its lags. Due to the observability provided by the sensor species, we can prove that generically there exists a local one-to-one map between the complete state of the ecosystem *x*(*t*) and the sensor space *Y_L_*(*t*) (Supplementary Note 1.5). This result implies that if an early-warning of a critical transition can be detected using all species in the ecosystem, then this transition can also be detected in the sensor space. To detect a transition in the sensor-space, we adopt early-warning methods developed to detect a critical slowing down from multi-dimensional time-series [24]. In particular, we used the correlation with lag 1 of the abundance of the sensor species, an increase of their variance, and an increase of either the maximum or minimum eigenvalues of the covariance matrix of the vector *Y_L_*(*τ*) ∈ ℝ^*p*(*N−p*)^ over a time-window *τ* ∈ [*t* − *T, t*] of size *T* > 0, denoted by λ_max,*C,T*_ and λ_min,*C,T*_, respectively. All of these indicators are early-warnings of critical transitions (Supplementary Note 2), being the increase in the variance one of the most widely used due to its simplicity.

We used the three-species ecosystem of Fig. 1 to illustrate the use of sensor species for detecting early warnings. In this toy ecosystem, we previously identified Y = {2} and Y = {3} as the minimal sets of sensor species, also concluding that species 1 cannot be a solo sensor species. We tested by simulation the effectivity of these sensor species for detecting early-warning signals of a critical transition, using as population dynamics of the ecosystem the GLV model (see see Supplementary Note 3.2 for details). To produce a critical transition in this ecosystem, we used a factor *μ* ≥ 0 to slowly and simultaneously decrease the interaction strength of species 1 with species 2 and 3, resembling an environmental change experimented by the ecosystem [25]. As *μ* approaches zero, the ecosystem experiments a critical transition at *μ* ≈ 0.1 (Fig. 2A). Using an increase in the variance of the sensor species abundance as an early-warning signal, we find that both sensor species produce early-warning signals ofthe critical transition (yellow and blue in Fig. 2B). By contrast, the non-sensor species cannot detect any signal of the critical transition before it happens (pink in Fig. 2B).

**FIG. 2:**
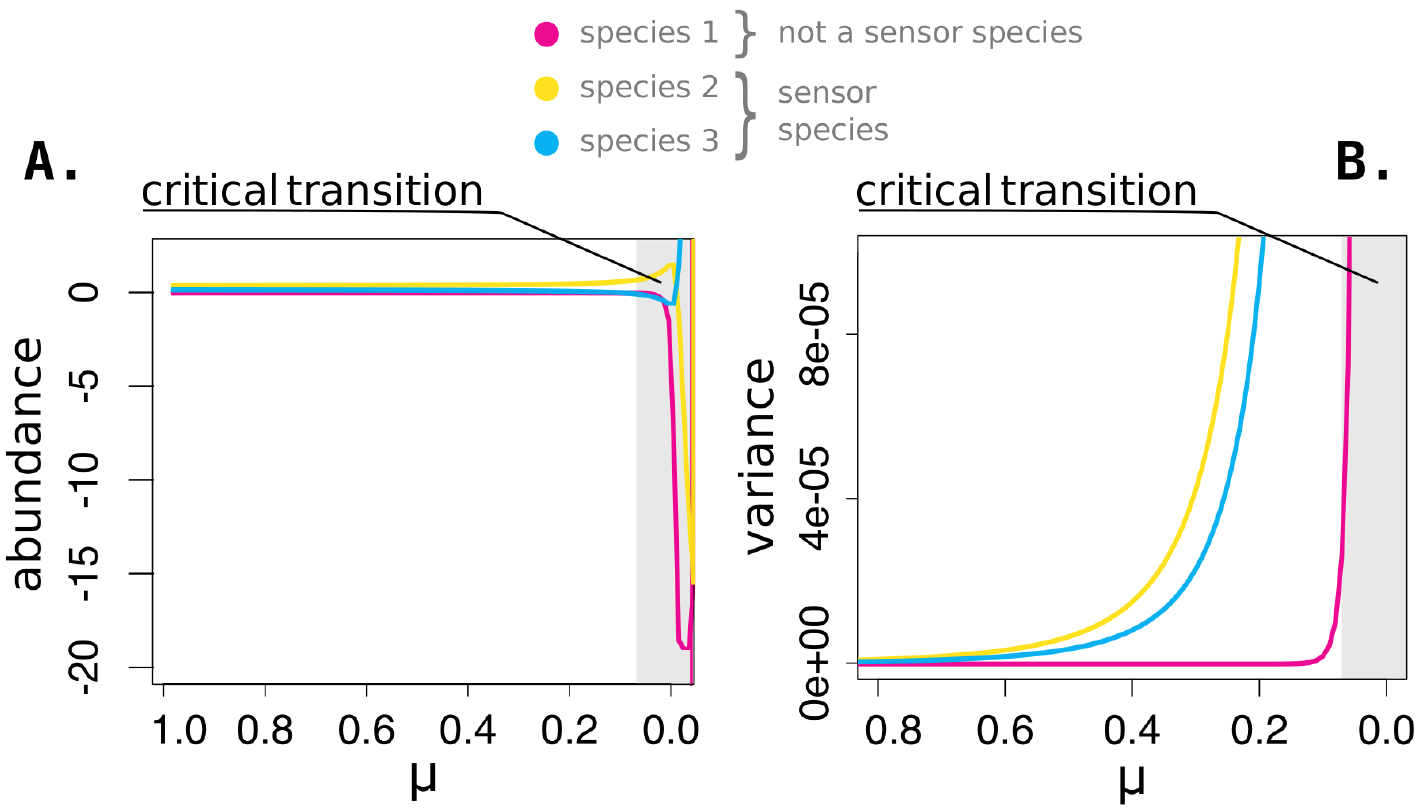
Using sensor species to detecting early-warning signals of critical transitions. **A.** To produce a critical transition in the ecosystem of Fig. 1A, the interaction strength of species 1 with species 2 and 3 was progressively decreased, simultaneously, using a factor that slowly changes its value from 1 to 0. The critical transitions occurs at *μ* 0.1. **B**. The variance in the abundance of the each species over a rolling window was used as an early-warning signal. The variance of the identified sensor species increases as the system approaches the critical transition (blue and yellow), providing an early warning of the transition. By contrast, the variance of the non-sensor species (pink) does not increase until the system is actually at the critical transition.

### Validation of sensor species in large ecosystems

To test the efficacy of sensor species for detecting early-warnings of critical transitions in large ecosystems, we simulated critical transitions in ecosystems with *N* = 500 species. We built those ecosystems using random ecological networks with two controlled parameters: their connectivity *c* ∈ [0,1] (i.e., proportion of edges with respect to a complete network), and the proportion of its edges that are bidirectional *ρ* ∈ [0,1]. For their population dynamics, we used Eq. (2) with GLV population dynamics with Holling’s type I or type II functional responses, given by the equations

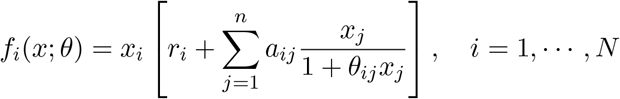

Above, the interaction matrix *A* = (*a_ij_*) ∈ ℝ^*N×N*^ is chosen as the adjacency matrix of the generated ecological network, and the parameters *θ_ij_* were chosen uniformly at random over the interval [0.01, 0.1]. The intrinsic growth rate vector *r* = (*r_i_*) ∈ ℝ^*N*^ was chosen to ensure that the initial state ofthe ecosystem is an equilibrium. Finally, we generated a critical transition by slowly decreasing the interaction strength of a randomly selected node, resembling an environmental change [25] (see details of the procedure in Supplementary Note 3.1). We identified a minimal set of sensor species for each ecosystem, and we use an increase in the variance of the sensor space over a rolling window as an early-warning indicator.

Regardless of the connectivity and proportion of bidirectional edges, early-warnings of the critical transitions were always detected in at least one sensor species (dotted lines in Fig. 3A). As comparison, we applied the same early-warning indicator to 10, 000 sets of randomly chosen species with the same size as the minimal set of sensor species that was identified. Early-warnings were detected only on a fraction of those random sets (solid lines in Fig. 3A), with the frequency of detection showing a high variability (Fig. S2 in Supplementary Note). The frequency of detection is lower for ecosystems with low connectivity or proportion of bidirectional edges. These two cases characterize condition under which the identified sets of sensor species provide significantly better information than measuring a randomly chosen species. As the connectivity or proportion of bidirectional edges increase, the frequency of detection in the random sets tends to increase. Indeed, in the limit cases when *c* = 1 or *ρ* =1 the frequency of detection of critical transitions using random sets of species is one, since in such cases any single species in the ecosystem is a minimal set of sensor species (Supplementary Note 3.1). The increased variability of the frequency of detection as *ρ* increases suggest that, just before the number of sensor species equals 1, the particular choice of species to measure becomes particularly relevant.

**FIG. 3:**
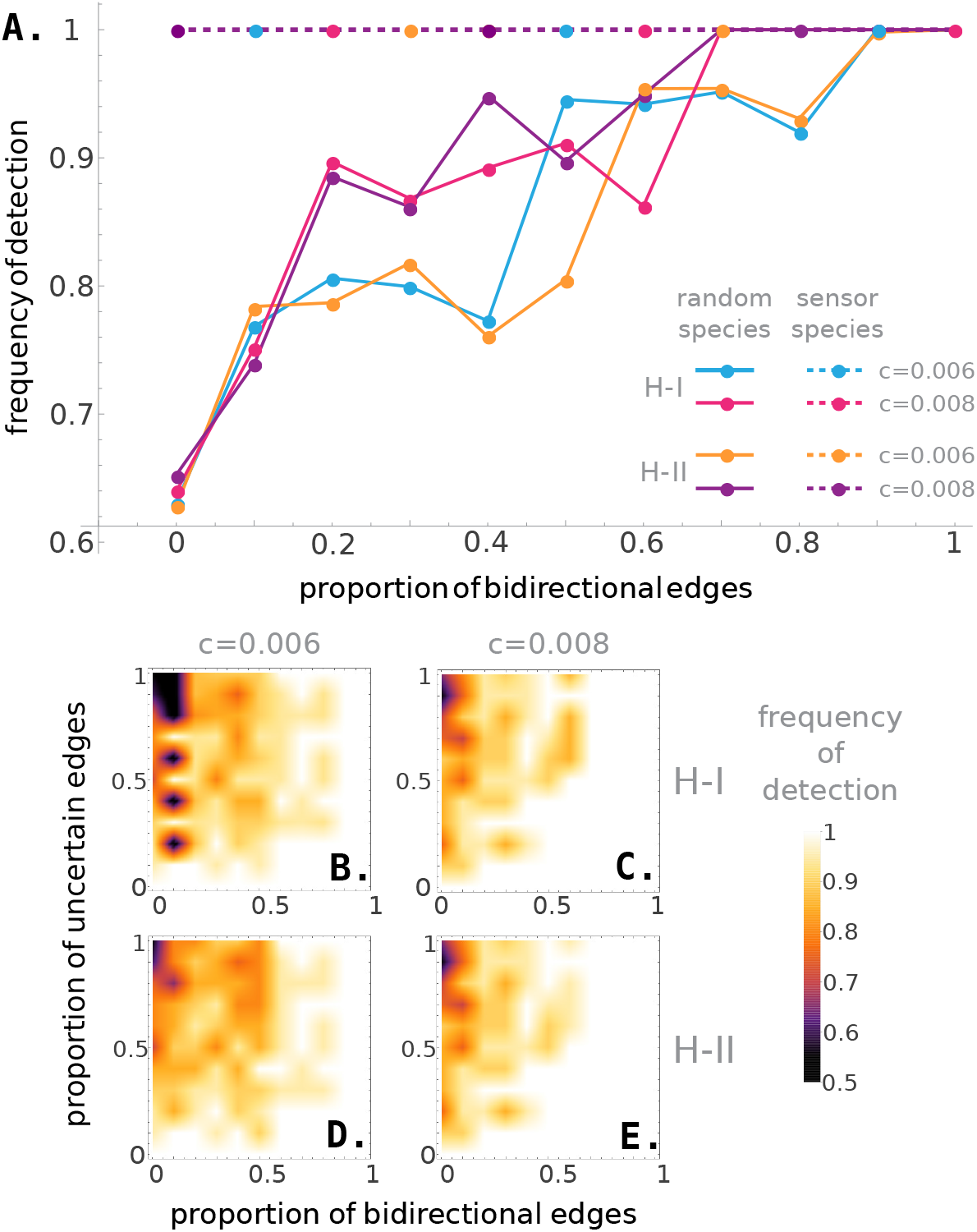
Sensor species detect critical transitions in large ecosystems. We generated ecosystems of *N* = 500 species having random ecological networks (with connectivity c and proportion of bidirectional edges *ρ*) and GLV population dynamics with functional responses of Holling’s type I and II. In each ecosystem, a critical transition was forced by disconnecting a randomly selected node and a minimal set of sensor species was identified. **A.** Mean frequency of detection of the critical transition in a minimal set of sensor species (solid lines), and in a random set of species with the same size (dotted lines), for a functional response of Holling’s type I (blue and purple), of Holling’s type II (red and orange), and connectivity *c* = 0.006 (blue and red) and *c* = 0.008 (purple and orange). The critical transition is always detected in the set of sensor species, and detected only in a fraction of cases in the set of random species. **B. and C.** Mean frequency of detection of the critical transition in a minimal set of sensor species with uncertainties in the network, for a functional response of Holling’s type I, and connectivity of *c* = 0.006 and *c* = 0.008 respectively. **D. and E.** Same as panels B. and C., with a functional response of Holling’s type II. The critical transition is detected in the majority of the cases despite uncertainties in the network.

We also tested the robustness of the minimal setofsensorspecies against errors in the ecological network used to identify it. We introduced errors in the ecological network by randomly rewiring a proportion *m* ∈ [0, 1] of its edges (e.g., *m* = 0.05 corresponds to a 5% error). We calculated a minimal set of sensor species from the rewired network, and then built early-warning indicators from this set of sensor species. Early-warnings of the critical transition are detected with high probability despite errors in the ecological network (Fig. 3B-E). More precisely, the transition was always detected in at least one species when *ρ* ≈ 1 and/or *m* ≈ 0. As *ρ* decreases or m increases, the frequency of detection lowers but it remains above 0.5 which means that in the worst cases the transition is detected in at least 50% of the analyzed ecosystems.

### Application to experimental data of a critical transition in a lake food-web

Finally, we applied our framework to detect early-warnings of a critical transition in a lake food-web using experimental data. The experimental data consists ofa three-year monitoring of two neighboring lakes [10], Peter and Paul, in which several types offishandplants cohabitate and interact. Adult predators were added on three occasions over two years to Peter Lake, inducing a critical transition to a state where predators dominate. By the end of the third year, this transition in the state of Peter Lake was considered complete. Paul Lake remained without manipulation (i.e., unperturbed) and, under the assumption that its population dynamics are similar to those of its neighbor lake, it was used as control for comparing the obtained early-warning signals. The ecological network underlying the lake food-web consists of adult and juvenile piscivores, planktivore fish, zooplankton, and phytoplankton [26] (Fig. 4A). Following our framework, we identified phytoplankton as the solo sensor species for this ecosystem (see Supplementary Note 3.3 for details). Note that the abundance of phytoplankton can be directly measured by monitoring chlorophyll levels and thus, our identified sensor species agrees with the experimental evidence showing that chlorophyll strongly responds to food web fluctuations [27, 28]. Early-warning signals in the sensor species are higher in Peter Lake than in Paul Lake (Fig. 4C), in agreement with previous analysis of this experimental data [10]. This result confirms that the identified sensor species provides robust early-warnings for the transition in this experimental ecosystem.

**FIG. 4:**
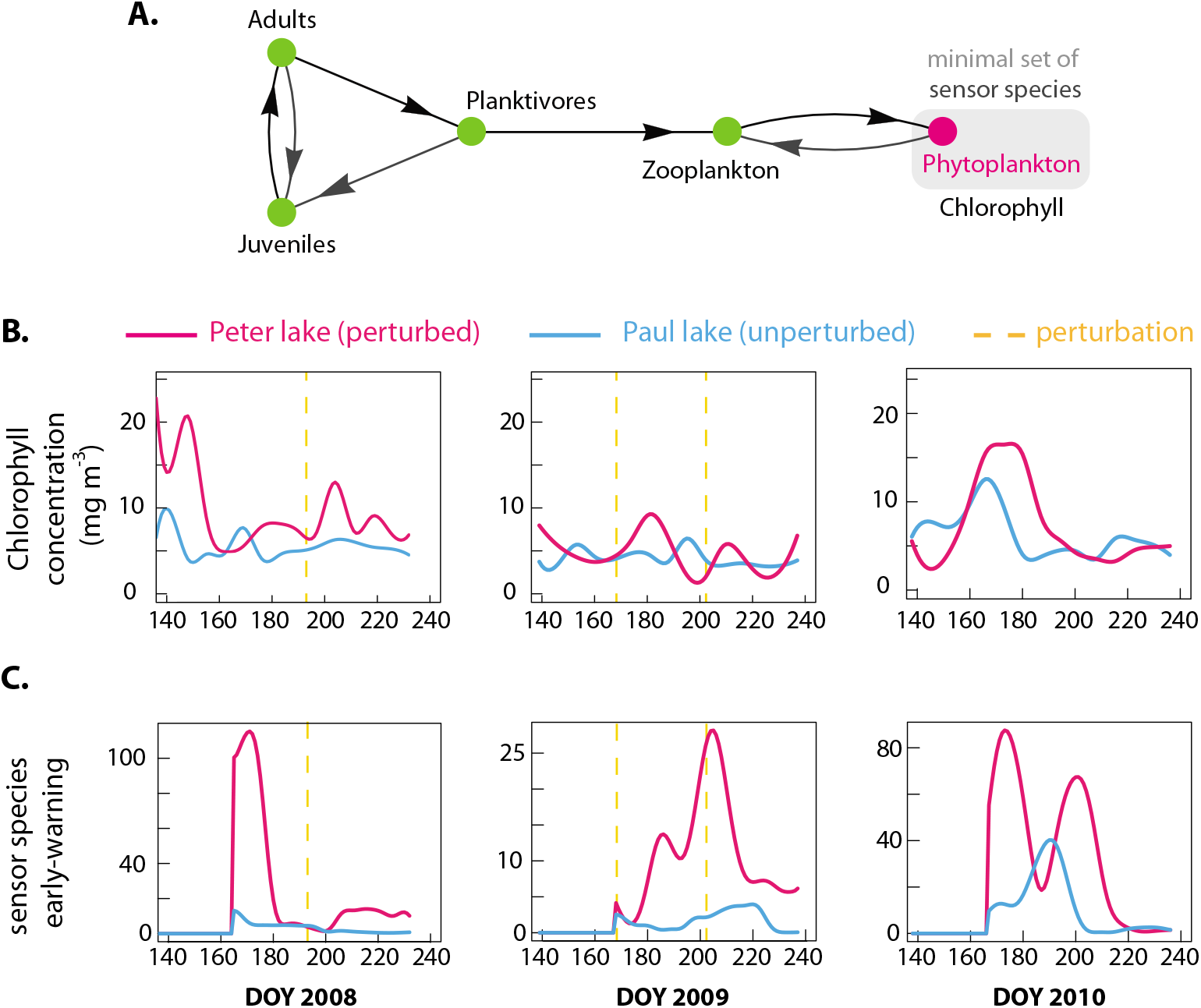
Predicting transitions in an experimental lake food web using sensor species. **A.** Ecological network of the lake food web, as in Ref. [26]. Black arrows mark a path covering all nodes, leading to a minimal set of sensor species containing only phytoplankton. Note that the phytoplankton can be measured by the concentration of chlorophyll. **B.** Chlorophyll concentration as a function of time for the years 2008, 2009 and 2010. Time instants when predators were added to Peter Lake (corresponding to perturbations) are marked by vertical yellow lines. **C.** Early-warning signals constructed from the sensor species using the maximum eigenvalue of the covariance matrix with *L* = 2 lags. High peaks in the sensor early-warnings are observed in Peter lake when predators are added to the ecosystem.

## DISCUSSION

Our mathematical formalism to identify a minimal set of species that needs to be measured to detect a critical transition in large complex ecosystems shows that these “sensor species” are generically characterized by the topology of the ecological network. In general, a minimal set of sensor species using our formalisms contains but is not equal to a minimal set of “sensor nodes” as defined using the graphical approach in Ref. [29]. Note also that the notion of sensor species introduced here is in general different from the existing notion of “indicator species”, which are species sensitive to certain environmental parameters such as fire or pollution [30]. In the formulation of our paper, the indicator species would be species that render certain parameters of the ecosystem observable. By contrast, the sensor species render the abundance of all the other species observable.

The advantage of identifying a minimal set of sensor species in an ecosystem of *N* species increases as the ratio *p/N* decreases, where *p* is the size of a minimal set of sensor species. We calculated this ratio for 116 empirical ecological networks from fisheries [31] having between 6 and 54 species, finding an average *p/N* ≈ 0.06 (Supplementary Note 3.4). This means that only 6% of the species needs to be measured in order to construct reliable early-warning signals, suggesting that sensor species could provide an efficient method to monitor and detect critical transitions in large real ecosystems. Indeed, the majority of these networks require a solo sensor species (Fig. S3 in Supplementary Note 3.4).

We end noting that the mathematical formalism we have developed can be applied without modifications for identifying “sensor components” when the ecosystem components are not species but ecological patches, functional groups, strains, or operational taxonomical units. Altogether, our results illustrate that the knowledge of the ecological network of complex ecosystems can be useful to detect imminent critical transitions before they occur. We have shown how the observability of complex networked systems [29] can be constructively used, allowing us to better monitor large ecosystems whose population dynamics remain highly uncertain and difficult to infer.

## Authors’ contributions

M.T.A. conceived and designed the project. A. A. and M.T.A. did the analytical calculations. A.A. did the numerical simulations and analyzed the empirical data. All authors analyzed the results. M.T.A. wrote the main manuscript with the help of A.A. All authors edited the manuscript.

## Acknowledgements

We thank the financial support of CONACyT, México. Fordecyt. We also thank Dr. Claire Jacquet and Prof. Stephen Carpenter for providing us with experimental data.

